# Electrophysiological effects of psilocybin co-administered with midazolam

**DOI:** 10.1101/2025.07.25.666887

**Authors:** Michael H. Sutherland, Christopher R. Nicholas, Richard C. Lennertz, Cody J. Wenthur, Bryan M. Krause, Christina J. Sauder, Brady A. Riedner, Richard F. Smith, Paul R. Hutson, Charles L. Raison, Matthew I. Banks

## Abstract

The serotonergic psychedelic psilocybin induces neural plasticity and profoundly alters consciousness. The benzodiazepine midazolam blunts neural plasticity and induces conscious sedation and amnesia at low doses. In our recent open label pilot study, we administered oral psilocybin (25 mg) along with intravenous midazolam at doses allowing a full psychedelic experience while blunting memory for the experience. We previously reported preliminary results from high density scalp electroencephalography (EEG) recorded during the dosing session. Here, we examined changes in EEG band power, normalized Lempel Ziv complexity (LZCn), and spectral exponent. We used linear mixed effects models that incorporated time and the subjective effects of midazolam and psilocybin, measured with the Observer’s Assessment of Arousal and Sedation (OAA/S) and selected items from the Altered States of Consciousness (ASC) questionnaire, respectively. At 15-30 mins, when midazolam (but likely not psilocybin) was at its targeted effect site concentration, we observed increased beta power and decreased spectral exponent. As the subjective effects of psilocybin commenced and over the next six hours, we observed increased LZCn and spectral exponent and decreased broadband power. OAA/S improved model fits for alpha power while ASC improved model fits for LZCn and spectral exponent. These data are further evidence that the effects of psilocybin are maintained in the presence of midazolam, supporting its utility in mechanistic studies of psilocybin’s therapeutic activity.

## Introduction

Psilocybin, a serotonergic psychedelic, and midazolam, a benzodiazepine sedative and positive allosteric modulator of GABA_A_ receptors, have divergent functional effects on the brain and subjective experience. Psilocybin induces neural plasticity and metaplasticity (de la Fuente Revenga et al., 2021; Hesselgrave et al., 2021; Nardou et al., 2023; Shao et al., 2021; Vargas et al., 2023) and induces a profoundly altered state of consciousness (Griffiths et al., 2011; Roseman et al., 2017), the ‘psychedelic experience’. Midazolam blunts neural plasticity and induces conscious amnesia at low doses (del Cerro et al., 1992; Evans & Viola-McCabe, 1996; Puig-Bosch et al., 2022; Veselis et al., 2009). Recently, we reported the results of a preregistered, open label pilot study (Nicholas et al., 2024) in which we co-administered psilocybin (25 mg) with midazolam at doses that induced mild to moderate sedation while blocking memory encoding. The combined administration produced a psychedelic experience comparable to psilocybin monotherapy, while memory for the experience was blunted in some cases. During the dosing session, we monitored brain activity with high density electroencephalography (EEG), sedation level using the Observer’s Assessment of Arousal and Sedation (OAA/S) (Chernik et al., 1990), and the intensity of the subjective experience using selected questions from the Altered States of Consciousness (ASC) questionnaire (Studerus et al., 2010). In our previous paper, we reported an acute reduction in EEG alpha power during psilocybin co-administration with midazolam; here, we report further exploratory analyses of EEG effects.

We focus on three EEG metrics that have previously been linked to changes in arousal state and modulation by psychedelics and anesthetic agents: frequency band power, spectral exponent, and normalized Lempel Ziv complexity (LZCn). Frequency band-specific changes in EEG signal power associate with changes in arousal and behavioral state (Buzsaki, 2006). For example, psilocybin has been shown to acutely induce a broadband decrease in signal power, with prominent reductions the alpha band (8-13 Hz) power that are often observed in the resting state (Muthukumaraswamy et al., 2013; Ort et al., 2023; Timmermann et al., 2019). Midazolam is also reported to reduce alpha band power, but, in contrast to serotonergic psychedelics, increases delta band (1-4 Hz) and beta band (20-30 Hz) power (Breimer et al., 1990; Hotz et al., 2000).

Spectral exponent describes the decay of the aperiodic component of EEG signal power with frequency. Computational models suggest that spectral exponent varies with excitation/inhibition (E/I) balance (Gao et al., 2017; Lombardi et al., 2017). Consistent with these models, spectral exponent decreases during sleep (Aamodt et al., 2022; Höhn et al., 2024; Kozhemiako et al., 2022; Lendner et al., 2020) and anesthesia (Colombo et al., 2019; Gao et al., 2017), both of which have been linked to increased GABAergic inhibition (Franks, 2008; Park et al., 2020; Tossell et al., 2023). Conversely, spectral exponent increases after administration of serotonergic psychedelics (Muthukumaraswamy & Liley, 2018), which promote neural excitability via 5-HT_2A_ receptor activation (Banks et al., 2021; Celada et al., 2013; Wallach et al., 2023).

LZC measures the compressibility of time-varying signals by counting the number of non-redundant binary patterns contained in the signal (Lempel & Ziv, 1976), and is associated with the repertoire of brain network states (Sarasso et al., 2021). Similar to spectral exponent, LZC reliably decreases with depth of sleep (Aamodt et al., 2021; Aamodt et al., 2022; Abásolo et al., 2015; Andrillon et al., 2016; Schartner, Pigorini, et al., 2017) and anesthesia (Hudetz et al., 2016; Maschke et al., 2024; Schartner et al., 2015; Wenzel et al., 2019; Zhang et al., 2001), and increases upon administration of serotonergic psychedelics (Mediano et al., 2024; Ort et al., 2023; Schartner, Carhart-Harris, et al., 2017; Timmermann et al., 2019).

Despite reliable findings when band power, LZC, and spectral exponent measures are applied to EEG collected from human participants administered either psychedelics or sedating agents in isolation, it remained unclear how these measures would behave when both psilocybin and midazolam were given concurrently. Thus, we measured band power, normalized LZC (LZCn), and spectral exponent at several timepoints throughout the combined psilocybin-midazolam dosing session. We sought to test: 1) whether psilocybin-induced changes in band power, LZCn, and spectral exponent observed in previous reports with psilocybin monotherapy remain with co-administration of midazolam, and 2) how band power, LZCn, and spectral exponent relate to arousal state and the subjective quality of the psychedelic experience during the dosing session, measured using OAA/S (Chernik et al., 1990) and the ASC (Studerus et al., 2010) respectively.

## Methods

### Ethics statement

Research protocols were approved by the University of Wisconsin Institutional Review Board (Protocol #2020-0085), and written informed consent was obtained from all participants. This study is registered on https://clinicaltrials.gov/study/NCT04842045.

### Dosing session and behavioral measures

Eight healthy participants were administered an oral 25 mg dose of psilocybin and IV midazolam doses targeting plasma concentrations of 25-95 mg/ml. The oral psilocybin and initial IV midazolam bolus were simultaneous. The initial midazolam bolus was chosen based on participant age and weight, with additional bolus adjustments over the next 3.5 hours to maintain mild to moderate sedation with memory impairment. Memory impairment was assessed with the California Verbal Learning Test (CVLT) [(Woods et al., 2006)], with the goal of achieving memory impairment such that CVLT scores were < 25%. Sedation was measured by scores on the OAA/S, with the goal of maintaining arousal level ≥ 3. OAA/S measurements were collected repeatedly between 5- and 210-minutes post-dosing. Subjective experience was assessed with 6 items from the ASC previously shown to predict antidepressant effect induced by psilocybin: profound thoughts, inventive ideas, inner peace, one with surroundings, time passing slowly/painful, colors in complete darkness (Roseman et al., 2017). ASC items were collected at 60-, 90-, 120-, 160-, 210-, and 360-minutes post-dosing, and the average score across the 6-items per time point was used for statistical modeling. For more details and a diagram outlining the dosing session, see (Nicholas et al., 2024).

### EEG and behavioral assessments

High-density EEG was collected continuously from 20-minutes pre-dosing to 360-minutes post-dosing. 256 channels were sampled at 500 Hz using a NetAmps 300 amplifier and NetStation software (Electrical Geodesics, Inc., Eugene, OR). From these data, 10 time points were chosen during the dosing session for EEG analysis: two baseline time points – music-on and music-off – and 9 post-dosing time points: 15-, 30-, 60-, 90-, 120-, 160-, 210-, and 360-minutes. Time points were chosen based on continuous (relatively) artifact free EEG and to match time points in which the OAA/S and ASC were assessed. All time points were periods with eyes-closed, and post-dosing time points were periods with music-on except for 30 and 120 minutes, which were music-off. In each EEG time point, 0.11-8.92 minutes of data immediately preceding the stated time was used for analysis (mean 4.04 minutes).

### Preprocessing of high-density EEG data

Preprocessing of EEG data was done using MATLAB and EEGLAB (Delorme & Makeig, 2004) software. Electrodes located on the face or neck were excluded leaving 185 channels per participant. First-order high-pass (single pole IIR, 0.1 Hz) and band-pass (2-way least squares FIR, 1–50 Hz) filters were applied. Bad channels and artifactual segments were rejected by visual inspection using EEGLAB. EEG data were denoised using Adaptive Mixture ICA (AMICA) in EEGLAB. Data were then visually inspected in component space and if artifactual segments remained, artifact rejection was performed again on the original data. If necessary, AMICA was run again on the original data with the additional noisy segments removed. Components labelled as eye, muscle, or channel artifact, and those labelled as neural but below a confidence threshold of 85% were removed. Rejected channels were then interpolated using spherical splines method and average referenced.

### Band power

Power spectral density (PSD) of the EEG signal was calculated using Welch’s method with 4-second nonoverlapping windows. The PSD was separated into canonical bands: delta 1-4 Hz, theta 4-8 Hz, alpha 8-14 Hz, beta 14-30 Hz, and gamma 30-50 Hz. Log_10_(power) was computed and averaged across frequencies within each band.

### Lempel Ziv complexity

Brain signal diversity was measured using time-domain LZC (Lempel & Ziv, 1976). For each channel, the EEG data were first split up into 4 second nonoverlapping segments (2000 samples each). The signal (each segment of each channel separately) was binarized using the median amplitude of the Hilbert transform and converted to a string of 0’s and 1’s. LZC of the binarized string was computed with the LZ76 algorithm (Lempel & Ziv, 1976). Starting from the beginning of the string, the algorithm iteratively compares a “search” substring to all prior substrings seen in the signal. If the search substring is already contained in earlier substring patterns, the search window extends. If the search substring is a unique pattern, the complexity count is increased by one. The search continues until the entire string has been assessed. The total complexity count was then normalized by dividing by the string’s theoretical upper bound, N/log_2_(N), where N is the length of the segmented signal. This returns a value, LZC, which typically falls between 0 and 1. To assess LZC above and beyond the influence of spectral power, a phase-shuffled surrogate signal was created by computing the Fourier transform of the original signal, randomly shuffling its phase, and computing the inverse Fourier transform to return to the time domain (Schartner et al., 2015). The same binarization and LZC computation procedure followed as above. This process was repeated 100 times, and LZC was normalized by the average phase-shuffled LZC, returning the measure LZCn.

### Spectral exponent

Spectral exponent was computed with the FOOOF algorithm (Donoghue et al., 2020). First, each 4 second PSD segment (frequency range 1 – 50 Hz) was fit with an estimation of its power law (aperiodic) component by taking the slope between power values at the first and last frequencies of the spectrum in log-log space. This fit was then subtracted from the PSD, leaving a flattened spectrum without the initial aperiodic component estimate. A threshold of the 2.5 percentile of the flattened spectrum as defined, and any power values falling below the threshold were considered part of the aperiodic component. A second fit was then performed on the PSD by taking the slope between all power values within the aperiodic component. This component was removed from the PSD, leaving a flattened spectrum thought to contain periodic components and noise. The maximum peak in the flattened spectrum was then found and fit with a Gaussian if it exceeded a defined threshold of 2.2 standard deviations of the flattened spectrum. The fit Gaussian was removed from the spectrum, and the algorithm began its peak search again. The search continued until no more peaks in the flattened spectrum exceed the threshold. The Gaussian fits were combined to create a periodic fit that was subtracted from the original PSD. A final aperiodic fit was then performed on the remaining PSD. Spectral exponent was taken as the negative of the final fit exponent, which corresponds to the slope of the full spectrum aperiodic estimation.

### Statistical modeling

Statistical analysis was performed using linear mixed effects models (*nlme* package) in R. To test the influence of time on band power, LZCn, and spectral exponent, models were run predicting each measure using time point (baseline and 15-, 30-, 60-, 90-, 120-, 130-, 160-, 210-, and 360-minutes post dosing) as a fixed factor and subject as a random effect. A continuous-time first order autoregressive covariance structure (corCAR1) was included to account for serial correlation among repeated observations derived from 4-second EEG segments within each subject across time. Music condition (on or off) was omitted from the model due to 1) a sensitivity analysis showing minimal effect of adjusting for music on model results (**Supplementary Figure 1**) and 2) insufficient data (only 30- and 120-minutes as music-off) to differentiate between the effect of music and time. Before model fitting, band power, LZCn, and spectral exponent were averaged across EEG channels per observation. Post model fitting, marginal mean band power, LZCn, and spectral exponent values for each time point and contrasts between time points were estimated using the *emmeans* package in R. Multivariate t distributions were used to correct for multiple comparisons between time points. To evaluate potential mediating pathways of drug effects, OAA/S and ASC were included in the models as time-varying mediators representing the behavioral effects of midazolam and subjective effects of psilocybin, respectively. Sequential likelihood ratio tests evaluated whether these mediational pathways significantly contributed to explaining variance in band power, LZCn, and spectral exponent within the time course of the dosing day.

### Data and code availability

Data and code necessary to reproduce figures and statistical analyses performed for this manuscript are available at (https://zenodo.org/records/17080375).

## Results

Administration of psilocybin with midazolam induced systematic changes in EEG power spectral density at various time points throughout dosing day (**Figure 1A**). Band- and time point-specific changes were identified using linear mixed effect models. Estimated marginal mean power values from each band were averaged across channels and plotted across time (**Figure 1B-F**). Time had a significant effect on all bands (all p < 0.001). For each band, post hoc comparisons were computed between post-dosing time points and baseline (**Supplementary Table 1**). Delta power significantly decreased relative to baseline at all time points between 60 – 360 minutes. Theta and alpha power decreased significantly relative to baseline at all post-dosing time points. Beta power increased significantly at 15 and 30 minutes and was significantly lower than baseline at 160-through 360-minutes. Gamma power decreased significantly relative to baseline at 30-, 60-, and 120-minutes. The influence of sedation (OAA/S) and subjective intensity (ASC) on band power was then assessed. First, OAA/S was added to the previous models to compare model fits with and without OAA/S using a likelihood ratio test. OAA/S improved model fits for alpha power only (p = 0.04). There was no effect of including ASC to the model for any bands. Next, linear mixed effect models were run predicting band power with time as a fixed effect and participant as a random effect (as stated above), but for each EEG electrode separately. Marginal mean power values for each channel are depicted on separate topo plots for each band and time point separately to visualize spatial changes in band power. (**Figure 2**).

**Figure 1.**
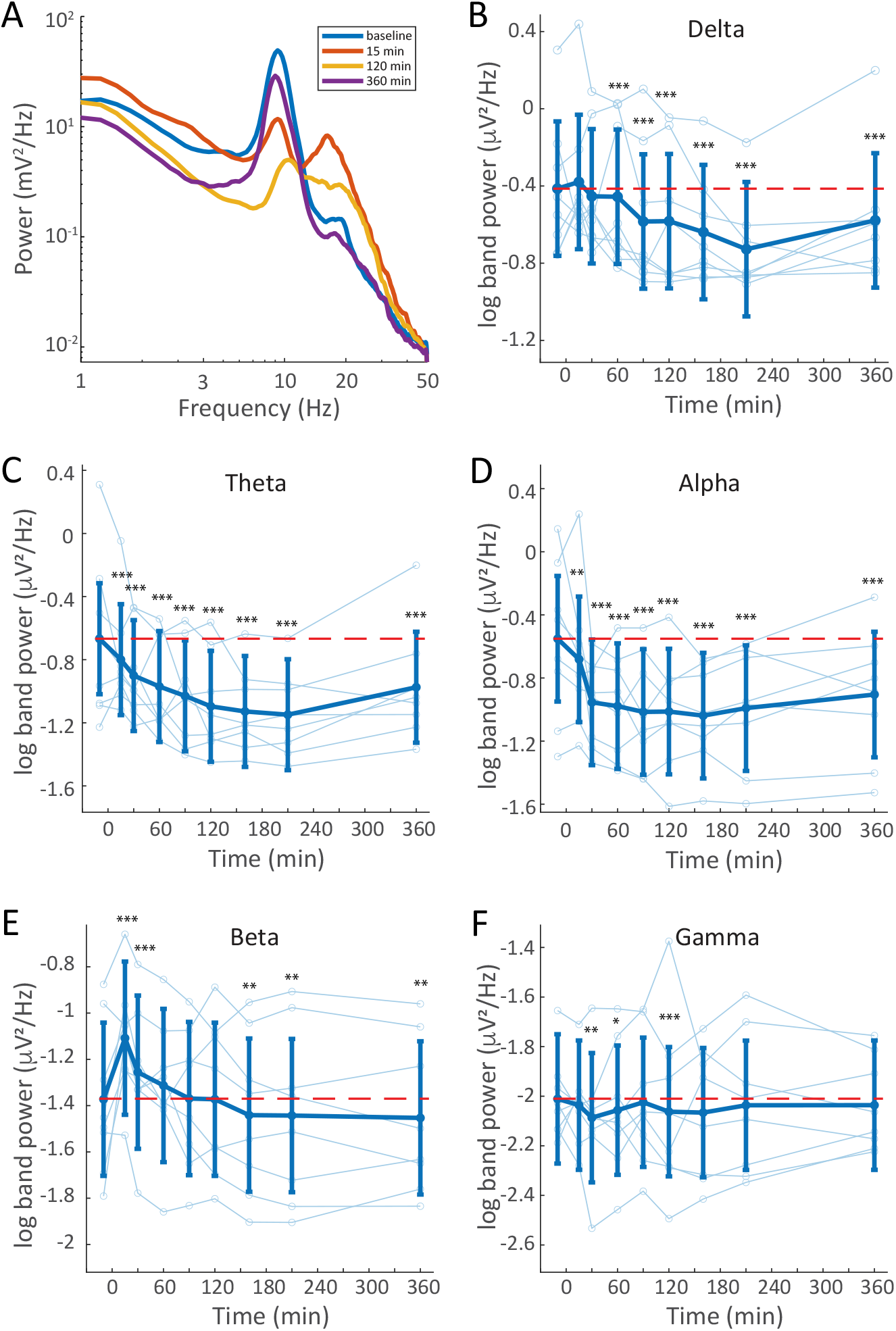
EEG power spectral density changes throughout dosing day. **A.** Lines represent eyes closed PSD from a single participant (PR004) at baseline, 15-, 120-, and 360-minutes post dosing. **B-F**. Time series plots of band power. Light blue lines and symbols show individual participant data; dark blue lines and symbols show model-derived marginal means and 95% confidence intervals. Panels (B) – (F) show delta (1-4 Hz), theta (4-8 Hz), alpha (8-14 Hz), beta (14-30 Hz), and gamma (30-50 Hz), respectively.

**Figure 2.**
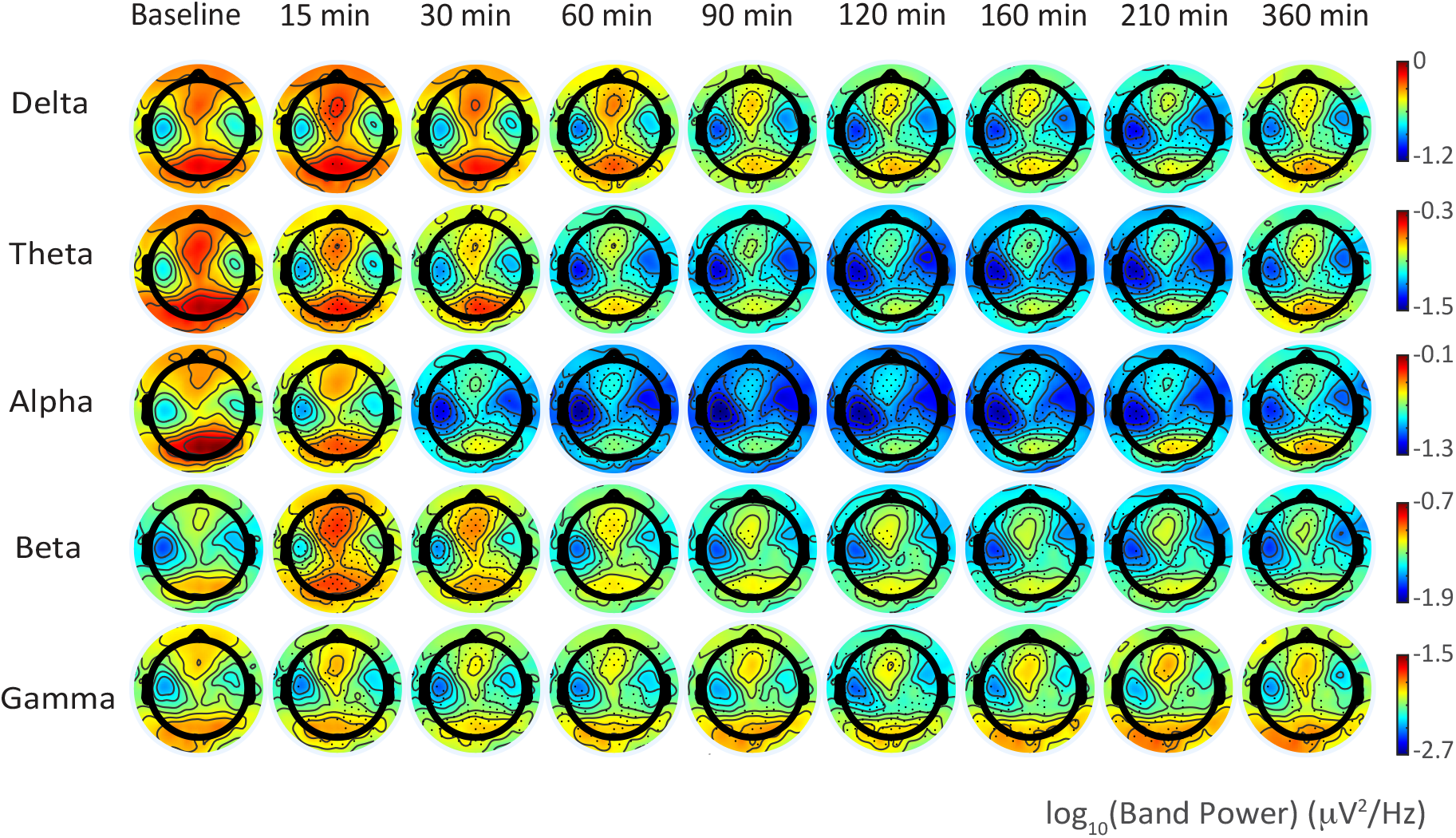
Topography of EEG power changes during administration of psilocybin with midazolam. Sensor-level topo plots for marginal mean band power derived from models fitting each channel separately. Each row is a frequency band; each column is a time point.

LZCn and spectral exponent consistently changed after administration of psilocybin with midazolam (**Figure 3**). A linear mixed effect model was run to assess the influence of time on LZCn, and the estimated marginal means were plotted across time (**Figure 3A**). Time had a significant effect on LZCn (p < 0.001). After running post-hoc comparisons between post-dosing time points and pre-dosing baseline, LZCn was found to be significantly increased from baseline at 60-through 210-minutes post-dosing, returning to baseline at 360 minutes (**Supplementary Table 2**). The effect of OAA/S and ASC on LZCn was subsequently assessed by including each measurement in the model as a fixed effect. When comparing models with each additional measurement to null models using a likelihood ratio test, OAA/S failed to improve the model fit for LZCn (p = 0.32). By contrast, ASC significantly improved the model fit for LZCn (p = 0.03). Model prediction of LZCn was plotted against the significant predictor ASC (**Figure 3B**), showing that increased ASC scores predicted increased LZCn.

**Figure 3.**
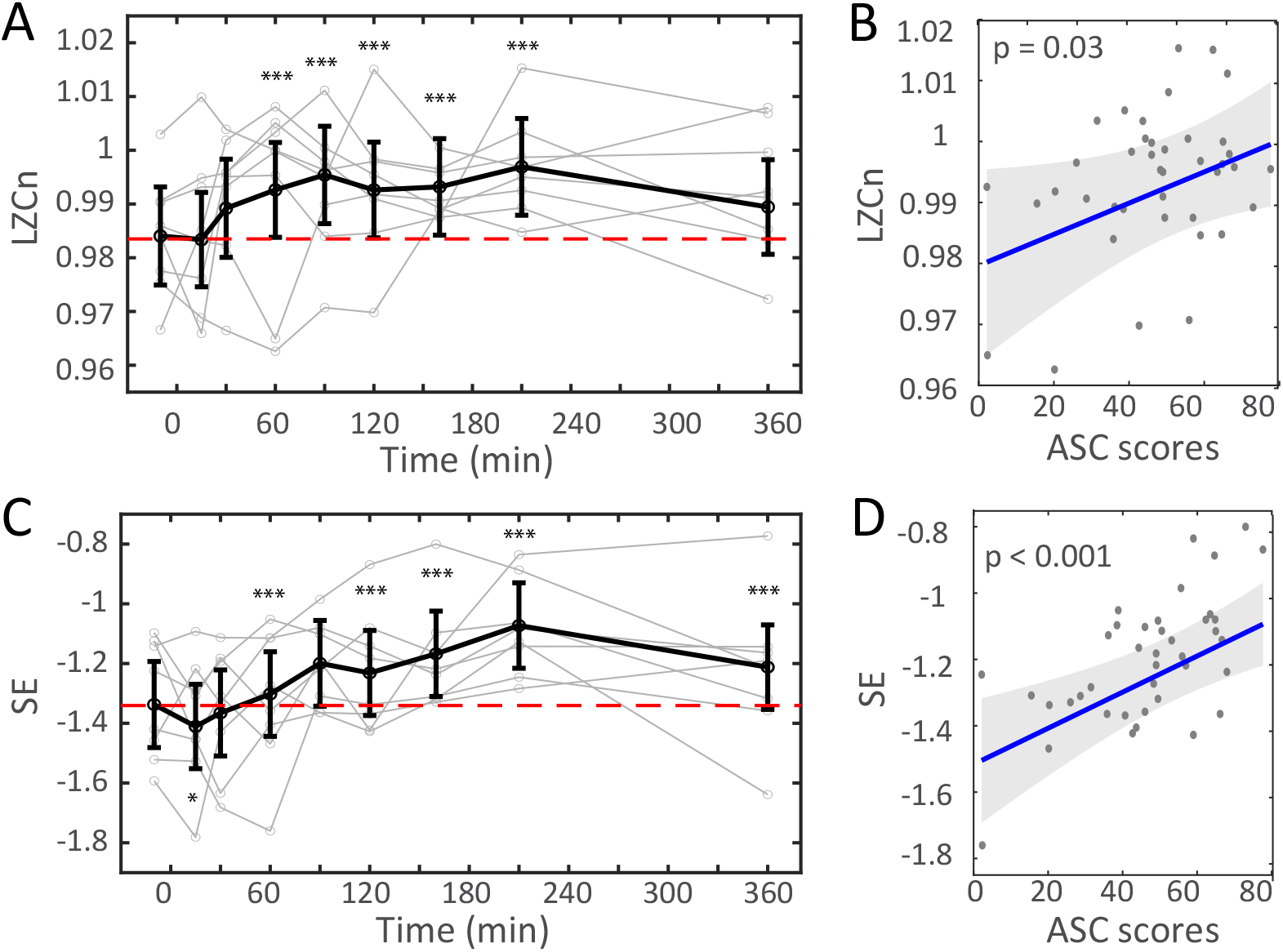
Complexity and spectral slope change systematically during administration of psilocybin with midazolam and depend on the subjective quality of the psychedelic experience. Shown are time series plots (A, C) of model-derived marginal means (black lines and symbols) and individual participant measurements (grey lines and symbols) for normalized Lempel-Ziv complexity (LZCn; **A**) and spectral exponent (**C**). Red dashed line represents baseline marginal mean value. (**B, D**) Model prediction of relationship between LZCn (**B**) and spectral exponent (**D**) with ASC scores.

A linear mixed effect model was then run to predict spectral exponent with time as a fixed effect and participant as random effect. Marginal means are plotted across time (**Figure 3C**), showing an increase in spectral exponent (i.e., a flatter spectrum) during the dosing session. Time (p < 0.001) had a significant effect on predicting spectral exponent. After running posthoc tests comparing post-dosing time points to baseline, a significant decrease from baseline was present at 15 minutes and significant increases were present at 90-through 360-minutes (**Supplementary Table 3**). After assessing the additional contribution of OAA/S and ASC to spectral exponent beyond time, OAA/S failed to improve model fit (p = 0.41) whereas ASC improved model fit for spectral exponent (p < 0.01). The relationship of the model-predicted spectral exponent versus the predictor ASC is depicted in **Figure 3D**, showing that increased ASC scores predicted increased spectral exponent.

## Discussion

### Summary

Changes in EEG spectral band power, LZC, and spectral exponent have previously been used to track changes in arousal, cognition, and consciousness, providing a link between neurophysiology and behavior (Aamodt et al., 2022; Buzsáki, 2006; Colombo et al., 2019; Lendner et al., 2020; Muthukumaraswamy et al., 2013; Muthukumaraswamy & Liley, 2018; Ort et al., 2023; Schartner et al., 2015; Schartner, Carhart-Harris, et al., 2017; Timmermann et al., 2019). Here, we show that the concurrent administration of psilocybin and midazolam in healthy individuals precedes changes in neural activity and subjective experience in a manner that is broadly consistent with these previous results. Broadband spectral power decreased during the dosing session, largely driven by decreases in delta, theta, and alpha bands. We observed a transient elevation of beta power that peaked at 15 minutes, when midazolam was estimated to have reached a steady plasma target concentration (Greenblatt et al., 1984) but before the subjective effects of psilocybin were apparent (Holze et al., 2023). LZCn increased during the dosing session before returning to baseline at 360 minutes, consistent with the time course of the acute effects of psilocybin. Spectral exponent decreased immediately post-dosing and then increased throughout the dosing session, remaining elevated at 360 minutes, broadly consistent with expected effects of both psilocybin and midazolam.

### Band power

At 15 minutes post-dosing, when midazolam but not psilocybin was expected to reach effect site concentrations sufficient to produce subjective effects (Crevoisier et al., 1983; Holze et al., 2023; Knoester et al., 2002), decreased theta and alpha power and increased beta power were observed, in agreement with prior studies on the effects of midazolam alone on EEG signal power (Breimer et al., 1990; Hotz et al., 2000). Once the effects of psilocybin became apparent, all bands decreased in power during the dosing session and all except gamma remained decreased at 360 minutes. Alpha power has been associated with the subjective effects of psilocybin, consistently decreasing according to the time course of acute drug effect (Muthukumaraswamy et al., 2013; Ort et al., 2023; Schartner, Carhart-Harris, et al., 2017; Timmermann et al., 2019). Here, alpha power reached its lowest values from 90 to160 minutes post-dosing, consistent with a typical time-window of peak subjective effects (Holze et al., 2023). Interestingly, however, including ASC scores in our model predicting alpha power did not significantly improve model fit, likely because only 1/6 of ASC items queried on dosing day related to visual processing (i.e., seeing colors with closed eyes).

### LZCn

LZCn has previously been shown to decrease with anesthetics and sleep depth and increase with serotonergic psychedelics (Aamodt et al., 2021; Aamodt et al., 2022; Abásolo et al., 2015; Andrillon et al., 2016; Hudetz et al., 2016; Maschke et al., 2024; Mediano et al., 2024; Ort et al., 2023; Sarasso et al., 2021; Schartner et al., 2015; Schartner, Carhart-Harris, et al., 2017; Schartner, Pigorini, et al., 2017; Timmermann et al., 2019; Wenzel et al., 2019; Zhang et al., 2001), likely reflecting changes in the temporal differentiation of underlying neural activity and the repertoire of network states. Given that deep sleep and some anesthetics are linked to enhanced GABAergic inhibition (Franks, 2008; Park et al., 2020; Tossell et al., 2023) and show decreased LZC, midazolam, through its GABA_A_ receptor modulation, is also expected to decrease LZC. This decrease in complexity with midazolam administration has been shown previously in spontaneous EEG (Murphy et al., 2023) and TMS-evoked EEG (Ferrarelli et al., 2010), although this is not always the case (Puglia et al., 2021). Serotonergic psychedelics, on the other hand, likely increase LZC through their purported 5-HT_2A_ receptor-dependent increases in neural excitability and network connectivity (Banks et al., 2021; Celada et al., 2013; Siegel et al., 2024; Wallach et al., 2023). Given the contrasting mechanisms of each drug, it was unclear what would transpire when co-administered. Our observation of an increase in LZCn suggests that the effects of psilocybin dominate over those of midazolam at these doses.

We also found that including ASC scores in our model improved model predictions of LZCn. Previously, it has been suggested that increases in brain signal complexity and entropy under administration of psychedelics reflects increased diversity of subjective states compared to typical waking consciousness (Carhart-Harris, 2018). The reported correlations between LZCn and aspects of the subjective experience of psychedelics (e.g. richness or intensity of experience) is consistent with this idea (Schartner, Carhart-Harris, et al., 2017; Timmermann et al., 2019). However, the link between LZC and the capacity for consciousness per se is less clear. For example, while LZC and LZCn were reported to increase during psilocybin administration previously (Ort et al., 2023) no change was observed in the Perturbational Complexity Index, arguably the most accurate measure of the capacity for consciousness to date (Casali et al., 2013). Furthermore, in a recent sleep study, LZC could not distinguish between conscious and unconscious sleepers (Aamodt et al., 2022). Moreover, in that study, LZC did not track aspects of the dream experience within the dream recall group, suggesting that LZC does not always reliably track subjective experience.

### Spectral exponent

Spectral exponent has previously been shown to decrease in sleep and anesthesia (Aamodt et al., 2022; Colombo et al., 2019; Gao et al., 2017; Höhn et al., 2024; Kozhemiako et al., 2022; Lendner et al., 2020). To our knowledge, only one study to date applied spectral exponent to data recorded during administration of serotonergic psychedelics, and it showed an increase from baseline resting state (Muthukumaraswamy & Liley, 2018). As with LZCn, including ASC scores in our model improved model predictions of spectral exponent. To our knowledge, this is the first report showing a significant relationship between aspects of subjective experience and spectral exponent (e.g. (Aamodt et al., 2022)) as opposed to the capacity for consciousness more broadly (Colombo et al., 2019; Zhao et al., 2025).

Spectral exponent has been linked to E/I balance in computational modeling (Gao et al., 2017; Lombardi et al., 2017), and spectral exponent changes in contexts of purported shifts in E/I balance, e.g., decreasing with shifts toward more inhibition. Our results are consistent with this idea, as midazolam’s GABAergic action is expected to shift this balance toward inhibition, with concomitant decreases in neural plasticity and memory. Spectral exponent, unlike LZCn, significantly decreased at 15 minutes post-dosing, where brain activity is more likely to reflect the effects of midazolam without contributions from psilocybin. However, our behavioral measure of midazolam sedation, OAA/S, remained an insignificant contributor in predicting spectral exponent, which may reflect the lower dose requirement for amnesia compared to sedation (Veselis et al., 2009). Additionally, unlike LZCn, spectral exponent remained significantly elevated at 360 minutes, consistent with elevated subjective effects of psilocybin observed in 4/8 participants at 360 minutes (Nicholas et al., 2024) and the significant contribution of ASC to predicting spectral exponent. Broadband spectral exponent has been shown to correlate with level of consciousness, specifically being higher in states in which conscious report is available (e.g., typical waking consciousness, dreaming, ketamine anesthesia) vs unavailable (e.g., under propofol anesthesia) (Colombo et al., 2019). We note, however, that more rigorous analysis is required to assess the direct electrophysiological contributors to spectral exponent.

### Limitations

There are several important limitations in this study that should be considered when interpreting results. The open-label design, small sample size, and substantial individual variability observed among participants limit generalizability. As the goal of the pilot study was finding an optimal midazolam dose to induce amnesia while allowing the psychedelic experience to unfold, different dosing regimens were used across participants, with the second 4 receiving higher doses than the first 4. This variability presented an opportunity, however, which we leveraged here to investigate the potential interactions between midazolam and psilocybin. Despite our limited sample size, we are confident that these results are not driven by outliers. However, we cannot claim that the results will generalize given a larger sample.

The LZCn and spectral exponent measures employed also have inherent limitations. LZCn requires binarization of the data that may result in information loss and reduced sensitivity. Moreover, LZC is influenced by changes in spectral power, especially in the low frequencies, though here we accounted for this by normalizing with a phase-shuffled surrogate signal. Low frequency power is also thought to contribute substantially to spectral exponent, especially when fitting broadband exponents. Spectral exponent is also subject to contamination by scalp muscle activity which may not be entirely removed during preprocessing, as suggested by a recent study investigating the effect on spectral exponent of changes in muscle tone during sleep (Kozhemiako et al., 2022). In the current study, we did not expect or observe evidence of dramatic changes in scalp muscle tone, but further investigation is required.

## Conclusions

Previous evidence suggests that psilocybin and midazolam exert opposing electrophysiological effects. We tested the effects of the co-administration of psilocybin and midazolam on the common EEG measures of band power, LZCn, and spectral exponent. We found decreased broadband power, increased LZCn, and increased spectral exponent throughout the dosing session, with LZCn and spectral exponent being at least partially dependent on the subjective quality of the psychedelic experience. Taken together, these results provide further support for the possibility that although midazolam blocked some aspects of memory during and after the acute experience, the subjective experience itself was not attenuated.

In our upcoming randomized, placebo-controlled follow-up study (RECAP2, https://clinicaltrials.gov/study/NCT06692192), we will systematically investigate the acute and post-acute electrophysiological effects of psilocybin and midazolam utilizing resting state and TMS-evoked EEG activity. We will link these neurophysiological effects to subjective experience and long-term behavioral outcomes of psilocybin.

## Declarations

### Authors’ contributions

Conceptualization: CRN, MIB, RCL, CJW, BMK, PRH, JDD, CLR, MHS. Funding acquisition: CLR. Data collection: CRN, RCL, RFS. Data curation: BAR, RFS, CJS, MHS. Data analysis: MHS, MIB, BMK. Visualization: MHS, MIB, BMK. Statistical analysis: MHS, MIB, BMK. Writing: MHS, MIB, CLR. Editing: MHS, MIB, BMK, CLR, RCL, CRN.

## Acknowledgements

We wish to thank Kristin Van Hyfte and Lucy Ptak for serving as research coordinators, Katie Chilton, MD for acting as a facilitator, and for the time and effort of our participants.

## Supplementary Tables

**Supplementary Table 1.** Contrasts between time points for EEG band power. All models were constructed as follows: Band power ∼ time + (1 | participant). Band power Estimates have units of log10(µV^2^/Hz).

| Contrast | Estimate | Lower CL | Upper CL | SE | df | t ratio | p |
| --- | --- | --- | --- | --- | --- | --- | --- |
| **Delta power** |  |  |  |  |  |  |  |
| 15 - base | 0.0356 | -0.0253 | 0.0965 | 0.0232 | 4481 | 1.5357 | 0.4781 |
| 30 - base | -0.0358 | -0.1044 | 0.0329 | 0.0261 | 4481 | -1.3702 | 0.5995 |
| 60 - base | -0.0963 | -0.1588 | -0.0338 | 0.0238 | 4481 | -4.0508 | 0.0004 |
| 90 - base | -0.2074 | -0.2743 | -0.1404 | 0.0255 | 4481 | -8.1424 | <0.0001 |
| 120 - base | -0.2356 | -0.2996 | -0.1716 | 0.0243 | 4481 | -9.684 | <0.0001 |
| 160 - base | -0.2693 | -0.3344 | -0.2041 | 0.0248 | 4481 | -10.8687 | <0.0001 |
| 210 - base | -0.344 | -0.4095 | -0.2785 | 0.0249 | 4481 | -13.8052 | <0.0001 |
| 360 - base | -0.1898 | -0.2512 | -0.1284 | 0.0234 | 4481 | -8.1244 | <0.0001 |
| **Theta power** |  |  |  |  |  |  |  |
| 15 - base | -0.1367 | -0.2046 | -0.0687 | 0.0259 | 4481 | -5.2853 | <0.0001 |
| 30 - base | -0.222 | -0.2985 | -0.1455 | 0.0291 | 4481 | -7.6213 | <0.0001 |
| 60 - base | -0.3624 | -0.4321 | -0.2927 | 0.0265 | 4481 | -13.6556 | <0.0001 |
| 90 - base | -0.434 | -0.5086 | -0.3593 | 0.0284 | 4481 | -15.2706 | <0.0001 |
| 120 - base | -0.508 | -0.5794 | -0.4367 | 0.0272 | 4481 | -18.7088 | <0.0001 |
| 160 - base | -0.4964 | -0.569 | -0.4238 | 0.0277 | 4481 | -17.9521 | <0.0001 |
| 210 - base | -0.5054 | -0.5785 | -0.4324 | 0.0278 | 4481 | -18.174 | <0.0001 |
| 360 - base | -0.3265 | -0.395 | -0.2579 | 0.0261 | 4481 | -12.5143 | <0.0001 |
| **Alpha power** |  |  |  |  |  |  |  |
| 15 - base | -0.1038 | -0.187 | -0.0206 | 0.0317 | 4481 | -3.2777 | 0.007 |
| 30 - base | -0.3838 | -0.4775 | -0.29 | 0.0357 | 4481 | -10.7588 | <0.0001 |
| 60 - base | -0.481 | -0.5664 | -0.3956 | 0.0325 | 4481 | -14.805 | <0.0001 |
| 90 - base | -0.5303 | -0.6218 | -0.4388 | 0.0348 | 4481 | -15.2381 | <0.0001 |
| 120 - base | -0.5094 | -0.5968 | -0.422 | 0.0332 | 4481 | -15.3199 | <0.0001 |
| 160 - base | -0.4872 | -0.5762 | -0.3982 | 0.0339 | 4481 | -14.3884 | <0.0001 |
| 210 - base | -0.432 | -0.5215 | -0.3424 | 0.0341 | 4481 | -12.6843 | <0.0001 |
| 360 - base | -0.3225 | -0.4065 | -0.2386 | 0.0319 | 4481 | -10.099 | <0.0001 |
| **Beta power** |  |  |  |  |  |  |  |
| 15 - base | 0.2309 | 0.1753 | 0.2865 | 0.0212 | 4481 | 10.9051 | <0.0001 |
| 30 - base | 0.1161 | 0.0535 | 0.1787 | 0.0238 | 4481 | 4.8717 | <0.0001 |
| 60 - base | 0.0236 | -0.0335 | 0.0807 | 0.0217 | 4481 | 1.0861 | 0.8081 |
| 90 - base | -0.0265 | -0.0876 | 0.0346 | 0.0233 | 4481 | -1.1407 | 0.7707 |
| 120 - base | -0.0353 | -0.0937 | 0.0231 | 0.0222 | 4481 | -1.5877 | 0.4428 |
| 160 - base | -0.0778 | -0.1372 | -0.0183 | 0.0226 | 4481 | -3.4352 | 0.004 |
| 210 - base | -0.0826 | -0.1424 | -0.0228 | 0.0228 | 4481 | -3.6274 | 0.0023 |
| 360 - base | -0.0748 | -0.1309 | -0.0187 | 0.0214 | 4481 | -3.5024 | 0.0032 |
| **Gamma power** |  |  |  |  |  |  |  |
| 15 - base | -0.0356 | -0.0947 | 0.0235 | 0.0225 | 4481 | -1.5831 | 0.4473 |
| 30 - base | -0.0844 | -0.1505 | -0.0182 | 0.0252 | 4481 | -3.3529 | 0.0055 |
| 60 - base | -0.0707 | -0.1312 | -0.0101 | 0.023 | 4481 | -3.0678 | 0.0139 |
| 90 - base | -0.0377 | -0.1023 | 0.027 | 0.0246 | 4481 | -1.5309 | 0.4837 |
| 120 - base | -0.1133 | -0.1752 | -0.0514 | 0.0235 | 4481 | -4.8115 | <0.0001 |
| 160 - base | -0.0561 | -0.1191 | 0.007 | 0.024 | 4481 | -2.3384 | 0.1034 |
| 210 - base | -0.0181 | -0.0815 | 0.0452 | 0.0241 | 4481 | -0.7521 | 0.9656 |
| 360 - base | -0.022 | -0.0816 | 0.0376 | 0.0227 | 4481 | -0.9717 | 0.8786 |

**Supplementary Table 2.** Contrasts between time points for LZCn ∼ time + (1 | participant).

| Contrast | Estimate | Lower CL | Upper CL | SE | df | t ratio | p |
| --- | --- | --- | --- | --- | --- | --- | --- |
| 15 - base | -0.0007 | -0.0062 | 0.0048 | 0.0021 | 4481 | -0.3202 | 1 |
| 30 - base | 0.0052 | -0.001 | 0.0114 | 0.0024 | 4481 | 2.1856 | 0.145 |
| 60 - base | 0.0086 | 0.0029 | 0.0142 | 0.0021 | 4481 | 4.0007 | <0.001 |
| 90 - base | 0.0114 | 0.0053 | 0.0174 | 0.0023 | 4481 | 4.9401 | <0.001 |
| 120 - base | 0.0086 | 0.0028 | 0.0143 | 0.0022 | 4481 | 3.9011 | <0.001 |
| 160 - base | 0.0091 | 0.0033 | 0.015 | 0.0022 | 4481 | 4.0871 | <0.001 |
| 210 - base | 0.0129 | 0.007 | 0.0188 | 0.0022 | 4481 | 5.7191 | <0.001 |
| 360 - base | 0.0054 | -0.0001 | 0.0109 | 0.0021 | 4481 | 2.5608 | 0.059 |

**Supplementary Table 3.** Contrasts between time points for spectral exponent ∼ time + (1 | participant).

| Contrast | Estimate | Lower CL | Upper CL | SE | df | t ratio | p |
| --- | --- | --- | --- | --- | --- | --- | --- |
| 15 - base | -0.0735 | -0.137 | -0.01 | 0.0242 | 4481 | -3.0418 | 0.015 |
| 30 - base | -0.0283 | -0.0997 | 0.0432 | 0.0272 | 4481 | -1.039 | 0.838 |
| 60 - base | 0.035 | -0.0301 | 0.1001 | 0.0248 | 4481 | 1.4118 | 0.569 |
| 90 - base | 0.138 | 0.0682 | 0.2077 | 0.0265 | 4481 | 5.1989 | <0.001 |
| 120 - base | 0.1057 | 0.039 | 0.1723 | 0.0254 | 4481 | 4.1669 | <0.001 |
| 160 - base | 0.1698 | 0.1019 | 0.2376 | 0.0258 | 4481 | 6.5736 | <0.001 |
| 210 - base | 0.2644 | 0.1962 | 0.3327 | 0.026 | 4481 | 10.1807 | <0.001 |
| 360 - base | 0.1252 | 0.0611 | 0.1892 | 0.0244 | 4481 | 5.1364 | <0.001 |

**Supplementary Figure 1.**
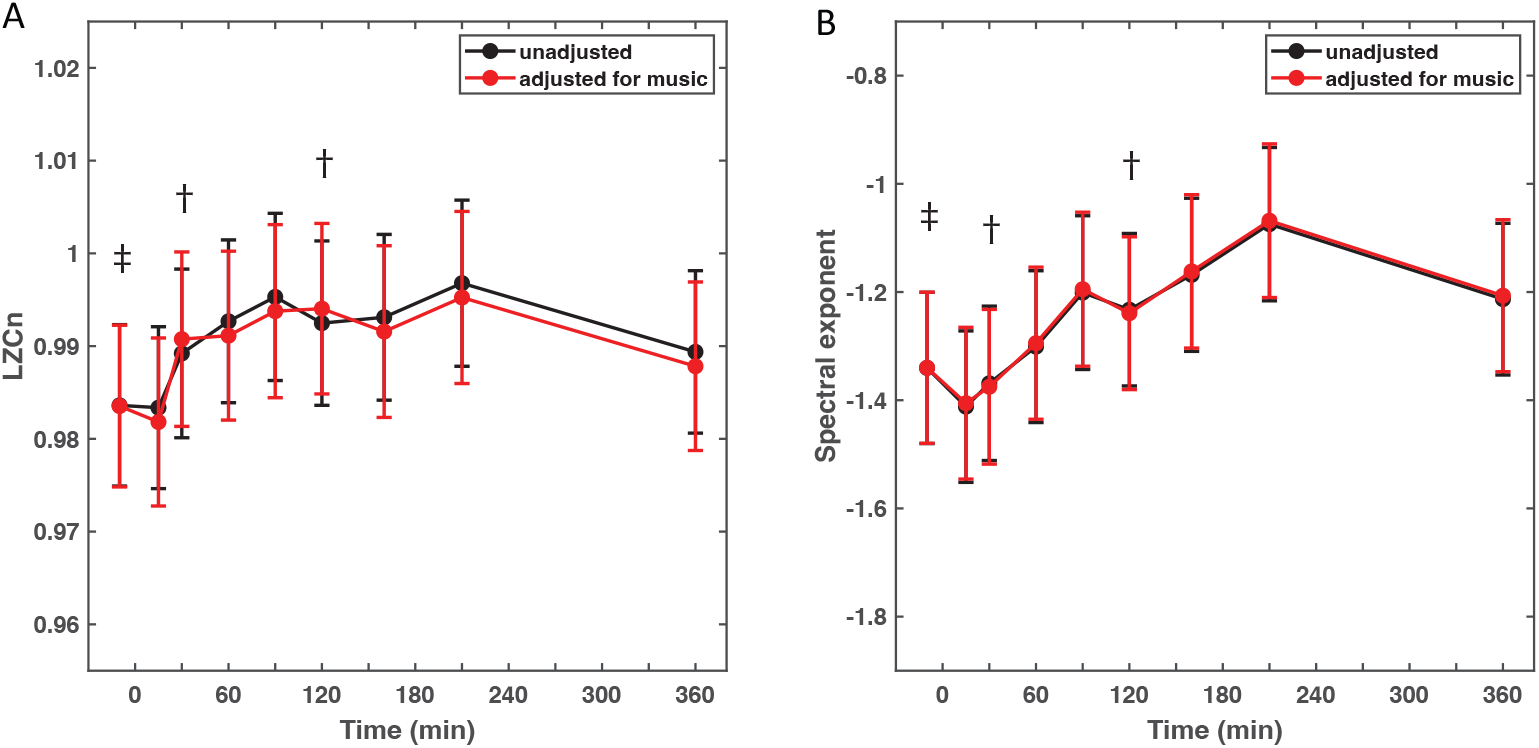
Complexity and spectral slope change little when adjusting for music. Shown are time series of marginal mean LZCn (**A**) and spectral exponent (**B**). Black lines represent marginal means derived from models run without adjusting for music (also shown in **Fig. 3A**,**C**), and red lines represent models adjusted for music (Measure ∼ time + music + (1 | participant). Symbol † represents time points with solely music-off data, and symbol ‡ signifies the time point with both music-on and music-off data.

